# Identification of novel basil downy mildew resistance genes using *de novo* comparative transcriptomics

**DOI:** 10.1101/2022.05.23.491563

**Authors:** Kelly S. Allen, Gregory A. DeIulio, Robert Pyne, Jacob Maman, Li Guo, Robert L. Wick, James Simon, Anne Gershenson, Li-Jun Ma

**Affiliations:** University of Massachusetts Amherst Department of Biochemistry and Molecular Biology, Amherst, MA, USA; Rutgers University Department of Plant Biology, New Brunswick, NJ, USA; University of Massachusetts Amherst Stockbridge School of Agriculture, Amherst, MA, USA

**Keywords:** *Ocimum basilicum* (sweet basil), *Peronospora belbahrii*, resistance, downy mildew, salicylic acid, transcriptomics

## Abstract

- Sweet basil (*Ocimum basilicum* L.) production is threatened by the oomycete pathogen *Peronospora belbahrii* causing basil downy mildew (BDM); BDM resistant cultivar ‘Mrihani’ (MRI) was identified in a germplasm screen, and fertile progeny were produced through a breeding program with BDM-susceptible ‘Newton’ (SB22), but the molecular mechanisms conferring resistance in MRI and progeny remained unknown
- Comparative transcriptomics was performed to identify candidate resistance genes and potential mechanisms for BDM resistance; RNA samples from BDM-infected MRI and SB22 plants were harvested at 4 time points during the first 3 days of infection to differentiate interactions in resistant and susceptible plants.
- Three categories of genes uniquely induced in resistant MRI upon pathogen challenge were identified: nucleotide-binding leucine rich repeat proteins (NLRs), multi-functional receptor-like kinases (RLKs), and secondary metabolic enzymes; validation of the top resistance candidate NLR gene confirmed its unique presence in MRI as well as in two of four resistant MRIxSB22 F_2_ progeny.
- In MRI, pathogen challenge also upregulated transcripts in the salicylic acid synthesis pathway, suggesting its role in BDM resistance, and demonstrating the application of using comparative transcriptomics to identify resistance genes and mechanisms in non-model crops for marker-assisted breeding approaches.

## INTRODUCTION

Basil (genus *Ocimum*) is a major herb crop with diverse species and cultivars possessing distinct phenotypes in plant size, leaf shape, aroma, and flavor (Vieira & Simon, 2006). Sweet basil (*Ocimum basilicum* L.) is the most popular basil and is cultivated for culinary use and essential oil production for applications including medicine, health care products, and food additives. In 2019, revenue generated in the US from sweet basil and other culinary herbs grown for dry processing and fresh market sales was estimated to be $165 million dollars, and other estimates have even valued the retail market above $300 million dollars (Wyenandt *et al*., 2015; *(dataset) USDA National Agricultural Statistics Service (2017).*)

Basil Downy Mildew (BDM), caused by the biotrophic oomycete *Peronospora belbahrii*, has become the most important disease posing a serious threat to global basil production since its introduction and movement in Europe in 2001 and the US in 2007 (Belbahri *et al*., 2005; Wyenandt *et al*., 2015). The pathogen enters plant tissue through open stomata or direct penetration of the upper cuticle, and colonizes leaf tissue intercellularly, causing characteristic symptoms of interveinal chlorosis several days after infection (Wyenandt *et al*., 2015; Cohen *et al*., 2017). Following sustained periods of high relative humidity (>85%), the pathogen produces sporangiophores bearing infective sporangia, which emerge through stomata and create a gray to dark-gray discoloration corresponding to interveinal chlorosis (Wyenandt *et al*., 2015; Cohen *et al*., 2017). The sporangia are aerially dispersed and cause polycyclic infections throughout large areas of production (Wyenandt *et al*., 2015; Cohen *et al*., 2017). After prolonged infection, the leaves desiccate and are abscised from the plant (Wyenandt *et al*., 2015; Cohen *et al*., 2017). Symptoms and signs of BDM disease render plants unfit for commercial sale (Wyenandt *et al*., 2015).

First reported in Uganda in 1930, BDM began to attract attention in 2001 when disease instances were increasingly reported across the world with reports from the Americas, Asia, and Europe (Garibaldi *et al*., 2004, 2005; McLeod *et al*., 2006; Khateri *et al*., 2007; Roberts *et al*., 2009; Ronco *et al*., 2009; Martínez de la Parte *et al*., 2010; Nagy & Horváth, 2011; Kanetis *et al*., 2014; Šafránková & Holková, 2014; Choi *et al*., 2016). BDM was first reported in the US in 2007 in Florida, and then in the Northeast the following year (Roberts *et al*., 2009; Wyenandt *et al*., 2015). As of 2021, BDM has been reported in 44 US states, including Hawaii and the District of Columbia (Wyenandt *et al*., 2015; McGrath, 2021, 2022a), threatening productivity in every growing region.

Chemical and cultural control methods to prevent infection and reduce disease spread include conventional fungicides (Homa *et al*., 2014; McGrath, 2020b), nocturnal fanning (Cohen & Ben-Naim, 2016), nocturnal illumination (Cohen *et al*., 2013), and daytime solar heating (Cohen & Rubin, 2015). However, these measures can be prohibitively time consuming and costly, necessitating the development of improved basil cultivars with BDM resistance. Initial screening of basil germplasm identified the *O. basilicum* ‘Mrihani’ (MRI) (Horizon Seed Co., Williams, OR) with significant resistance to BDM (Pyne *et al*., 2015). This cultivar has a unique anise/fennel aroma and flavor profile, and a distinctive phenotype as compared to other *O. basilicum* cultivars selected for culinary use. For example, the MRI leaves are smaller than other basil cultivars and have serrated leaves compared to the large downward cupped smooth leaves of commercial sweet basil (Figure 1A). Most importantly, the taste and smell of MRI due to its unique methyl chavicol chemotype differs considerably from eugenol-enriched sweet basil (Pyne *et al*., 2015). This cultivar was used in a parental cross with BDM-susceptible and the Fusarium-resistant ‘Newton’ (also referred to as Rutgers breeding line SB22), and successfully produced fertile offspring. From the six-generation breeding design, four BDM resistant cultivars (‘Devotion’, ‘Obsession’, ‘Passion’, and ‘Thunderstruck’) were selected from the backcrossed population progeny with improved downy mildew resistance and desirable phenotypes and chemotypes (Simon *et al*., 2018).

**Figure 1:**
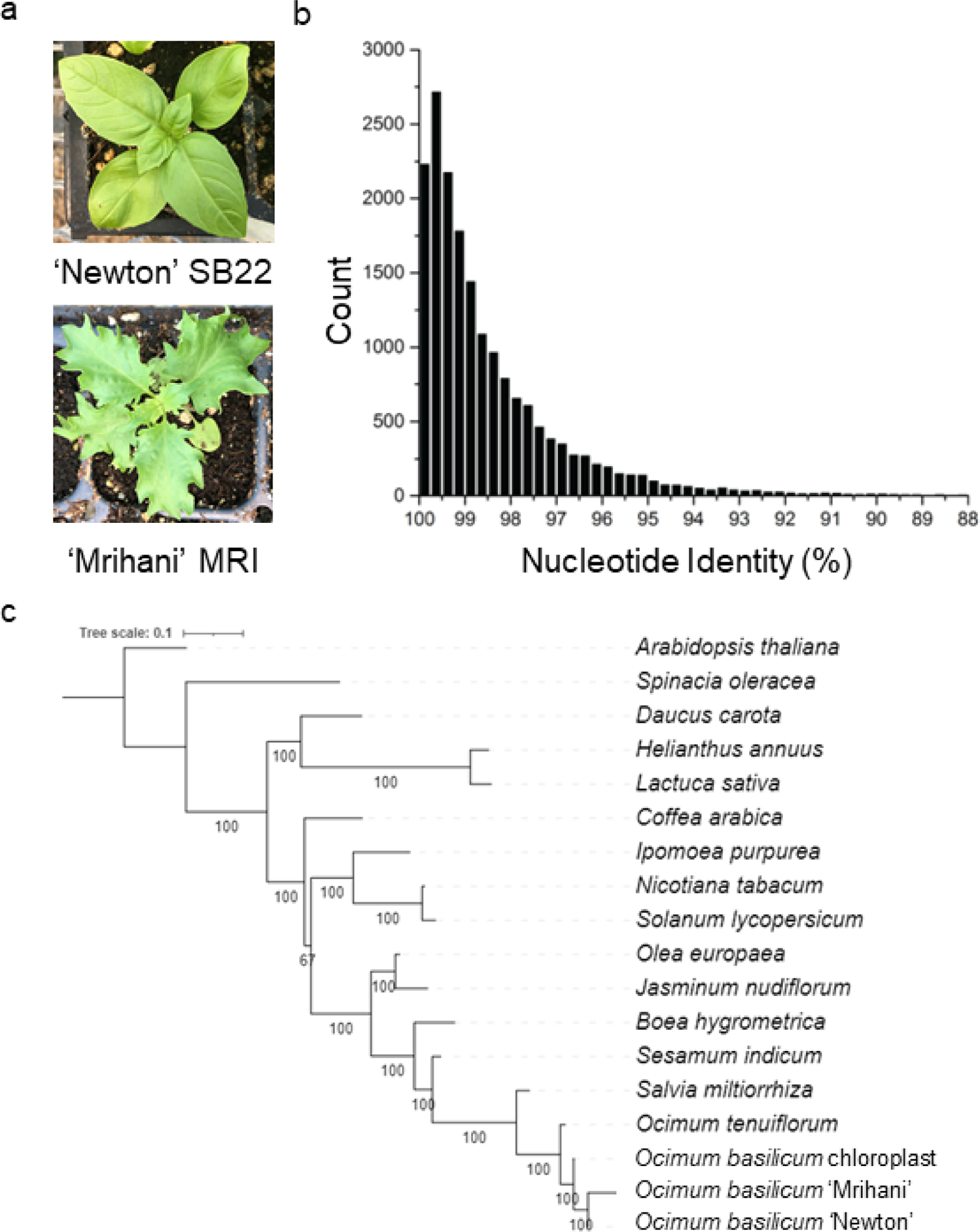
Basil cultivar phenotypic and genetic diversity. (a) BDM susceptible ‘Newton’ (SB22) and resistant ‘Mrihani’ (MRI) cultivars. (b) Bi-directional blast hit among 20,943 genes shared between the SB22 and MRI water references. The nucleotide identity for each top BLAST hit is graphed here with a bin size of 0.25%. (c) Multigene phylogeny of chloroplast genome orthologs across members of the asterid clade.

Screening of the full-sibling family offspring of the MRI x SB22 cross revealed additive and dominant gene effects of the MRI-conferred resistance, leading to the hypothesis that dominant alleles are involved in resistance (Pyne *et al*., 2015). A quantitative trait locus (QTL) analysis of the F_2_ mapping population from the cross between MRI and SB22 identified a single locus *(dm11.1)* that accounts for 20%-28% of the variance observed among the F_2_ population (Pyne *et al*., 2017). In addition, this study also identified two minor loci *(dm9.1 and dm14.1)* that respectively contributed 5-16% and 4-18% of the F_2_ population’s phenotypic variation. Resistance (R) genes involved in quantitative disease resistance map to QTLs (Nelson *et al*., 2018), and the results of the linkage mapping performed in the ‘MRI’ X SB22 F2 mapping population suggest that these loci may contain genes conferring quantitative disease resistance in ‘MRI’. Without genome assemblies of the basil cultivars, the underlying causative genes remained unknown.

A comparative transcriptomic analysis was designed to identify unique genes involved in the interactions of the resistant cultivar MRI and the susceptible commercial cultivar SB22 with the pathogen *P. belbahrii.* RNA was extracted at 12, 24, 48, and 72 hours post inoculation (hpi) representing roughly germination, penetration, and intercellular growth stages. Global transcription expression profiles identified three categories of genes uniquely induced in the MRI cultivar upon pathogen challenge, including R genes encoding nucleotide-binding leucine rich repeat proteins (NLRs), receptor-like kinases (RLKs) that sense conserved microbe-associated molecular patterns, and secondary metabolic enzymes. Validation of the top candidate resistance NLR protein-encoding gene confirmed its unique presence in the MRI cultivar as well as two out of the four resistance hybrids. Unique upregulation of the salicylic acid synthesis pathway in MRI suggests the perturbation of this important hormone signaling pathway in conferring BDM resistance.

## MATERIALS AND METHODS

### Sample preparation

Inbred *O. basilicum* genotypes SB22 (*P. belbahrii* susceptible) and MRI (*P. belbahrii* resistant) plants were grown from seed. Previously infected sweet basil leaves with fresh *P. belbahrii* sporulation were harvested and agitated for 5 minutes in sterile distilled water (diH_2_0). The inoculum mixture was filtered with 40 μm nylon mesh. A 1 mL subsample from the filtered inoculum was pipetted into an Eppendorf tube and frozen at −80°C to serve as a pathogen control. The remaining inoculum was centrifuged at 1,000g for 1 min and diH_2_0 decanted. The resulting sporangia pellet was resuspended in diH_2_O, and the inoculum concentration was adjusted to 1 x 10^5^ sporangia/mL. Four-to-six-week-old MRI and SB22 plants were spray-inoculated at the 6-leaf (3 true leaf set) growth stage with approximately 1 mL/leaf and plants were incubated at 100% relative humidity for 24 hours. A set of MRI and SB22 plants were sprayed with diH2O in triplicate to serve as the mock inoculated control.

Four disks per true leaf were sampled from both genotypes at 12, 24, 48 and 72 hpi and immediately flash frozen in liquid nitrogen. The water control leaves were harvested at 12 hpi only. Total RNA was extracted from freshly ground tissue using the Spectrum^™^ Plant Total RNA Kit (Sigma Aldrich). RNA samples were used to generate sequence libraries using a library prep kit from New England Biolabs (NEB #E7530). Paired-end sequence reads of 75 bp were generated at the TUFTs genomic center at the Tufts University School of Medicine using the Hi-Seq Illumina platform. Higher coverage analyses were specifically designed for inoculated samples that contain both the pathogen and the host due to the increased complexity of these samples, allowing for further study of pathogen expression.

### Generating transcript assemblies and FPKM expression

FASTQC version 0.11.5 (Andrews, 2010) was used to assess average read quality. Paired-end reads (fastq files) were provided to Trinity version 2.4.0 and assembled using default parameters (Grabherr *et al*., 2011). Datasets were assembled including single sets using either all MRI datasets and the sporangia control (MRI Combined Assembly) or all SB22 datasets including the sporangia control (SB22 Combined Assembly). Separately assembled control data provided organism specific databases of genes and transcripts.

The resulting output files served as the references for expression quantification. RSEM version 1.2.29 (Li & Dewey, 2011) and bowtie version 1.0.0 (Langmead *et al*., 2009) were used to calculate FPKM (Fragments per Kilobase exon per Million mapped reads) values for assembled contigs while tracking replicate information. RNAseq datasets from both MRI and SB22 were mapped to the infected MRI Combined Assembly to standardize the reference which allowed us to use previously generated gene annotations and to cluster genes from both cultivars together. In all cases the standard settings were used for assembly and transcript quantification. Additionally, edgeR (Robinson *et al*., 2010) was used to calculate differential gene expression. Expression data from both MRI and SB22 data mapped to the MRI Combined Assembly using Trinity and edgeR was used to assess differential expression between all timepoints within a single cultivar. Trinity DEG output data was filtered for genes with a p-value less than 0.05 and FDR less than 0.01.

### Reciprocal BLAST hits and reference gene phylogeny

To explore the overall sequence conservation between MRI and SB22, we performed a reciprocal BLAST using all sequences within the MRI and SB22 water control assemblies. Briefly, all MRI sequences were compared to the SB22 transcriptome, and all SB22 sequences were compared to the MRI transcriptome. The hit with the highest BLAST score for each gene was chosen. Results were compared and pairs of top scoring genes were considered reciprocal best BLAST hits (i.e., MRI gene X BLASTs to SB22 gene Z, and SB22 gene Z BLASTs to MRI gene X).

Sequence conservation between MRI and SB22 was further assessed by performing a phylogenetic analysis using 9 protein-coding chloroplast genome genes based on a prior analysis (Rastogi *et al*., 2015). The MRI and SB22 chloroplast genes were identified using BLAST against the MRI and SB22 water control assemblies, and the top hits with the highest bit scores were chosen and translated into coding sequences.

Sequence alignment of the MRI and SB22 coding sequences was performed against sequences from 14 asterid lineage plants downloaded from NCBI Organelle Genome Resources database, with *Spinacia oleracea* L. and *Arabidopsis thaliana* L. set as outgroups (Rastogi *et al*., 2015). The gene sequences were aligned using MAFFT (Madeira *et al*., 2019), and the tree was generated using IQ-TREE (Minh *et al*., 2020).

### Sequence Translation and Annotation

We generated a database of sequence annotations for MRI genes. All MRI genes with an expression of FPKM >1 in at least one time point were chosen and the longest transcript associated with that gene was compared to the NCBI non-redundant database using cloud BLAST through Blast2GO. Annotations were saved as a searchable database in text format. Genes were filtered by taxonomic hit to verify their species of origin as needed.

To facilitate easier searches for gene families of interest, we translated the longest nucleotide sequence associated with each gene into six-frame translated protein sequences using EMBOSS, searching for only those translated sequences between START and STOP codons longer than 30 amino acids. In many cases to verify protein domain structure, the nucleotide or protein sequence was analyzed using either the NCBI conserved domain finder (Marchler-Bauer *et al*., 2015), PFAM (Finn *et al*., 2016), or InterProScan (Jones *et al*., 2014). Sequences were aligned using MEGA 6 for visual inspection (Tamura *et al*., 2013).

### Expression clustering toward candidate resistance gene identification

Genes were clustered using Trinity version 2.2.0 based on read counts following the steps outlined in the Trinity manual (Robinson *et al*., 2010). Expression data generated by mapping all datasets to the MRI Combined Assembly were used for clustering. A matrix of gene expression at all timepoints and replicates was used to define clusters with the edgeR function associated with Trinity, using p=50 and p=20 (a grouping parameter for cluster creation, with higher numbers forming larger and broader clusters). The resulting clusters, available in pdf format, were visually examined for clusters which displayed the target expression profile.

### BLAST search for secondary metabolite enzyme and resistance genes

To analyze MRI unique gene families and defense hormone signaling genes, we performed BLASTp search against MRI and SB22 translated nucleotide sequences using *A. thaliana*, or in some cases sweet basil, protein sequences retrieved from NCBI. Generally, the hit with the highest bit score was chosen as the top hit for each sequence. In cases of short alignment length or low sequence identity, the recovered MRI or SB22 hit was compared to the green plant database on NCBI. BLAST version 2.2.22 was used in all cases to compare protein sequences (Altschul *et al*., 1990) at the Massachusetts green-energy high performance computing center (MGHPCC).

### PCR screen of parent and cultivar genomic DNA for unique genes

Genomic DNA was prepared from approximately 80mg of newly emerging leaf tissue of MRI, SB22, ‘Devotion’, ‘Obsession’, ‘Passion’ and ‘Thunderstruck’ cultivars using the E.Z.N.A. SP Plant DNA Kit (Omega BioTek, Norcross, GA) (Pyne *et al*., 2017). Primers amplifying transcript sequences were designed for MRI and SB22 shared genes as well as MRI unique genes including the comp160460c0 transcript, and were ordered from IDT (Coralville, IA). The primers were either external primers designed to amplify the whole gene (MRI_134-F 5’-CCGAGAAAATCGATCTAGAGAG-3’, MRI_2869-R 5’-CTAGCTTGATCTTTTAATTGGTGGAAAAAT-3’) or internal primers for specific regions of interest (Supporting Information Table S1). Primers amplifying a 198bp fragment of the *O. basilicum* Actin gene (ObActin_2-F 5’-GTTATGCACTTCCCCATGCT-3’, ObActin_2-R 5’-GAGCTGTTCTTTGCGGTCTC-3’) were used in positive control reactions for all cultivars. PCR was performed with Q5 High-Fidelity DNA Polymerase (New England Biolabs, Ipswich, MA) using manufacturer-recommended cycling conditions with a 30 second denaturation cycle to ensure full denaturation of genomic DNA, a 30 second extension time to amplify the 198bp ObActin region and a 120 second extension time to amplify the ∼3.5kb comp160460-encoded gene on a Mastercycler proS (Eppendorf, Hamburg, Germany). Water was used as a negative control template in all reaction sets. Amplicons were visualized on 1.5% agarose gels stained with SYBR Safe DNA Gel Stain (Invitrogen, Waltham, MA), and imaged under UV light.

### NLR allele analysis

Successfully amplified comp160460c0 products from MRI, ‘Devotion’ and ‘Obsession’ were purified using a QIAquick PCR purification kit (QIAGEN, Hilden, Germany) and single amplicon copies were ligated into the pMiniT 2.0 vector using the NEB PCR Cloning Kit (New England Biolabs, Ipswich, MA). Individual clones were selected and confirmed via colony PCR using Q5 High-Fidelity DNA Polymerase (New England Biolabs, Ipswich, MA) with primers 1BF and 8R which were designed to amplify the coiled coil and NB-ARC domain coding sequences of MRI-R1. Plasmid DNA was prepared from confirmed clones using the Zyppy Plasmid Miniprep Kit (Zymo Research, Irvine, CA).

Individual comp160460c0 clones from MRI, ‘Devotion’ and ‘Obsession’ were sequenced using NEB PCR Cloning Kit Cloning Analysis Forward and Reverse primers to flank the insert, as well as internal primers (Supporting Information Table S1). Twelve clones were sequenced from MRI, and six clones were sequenced from both ‘Devotion’ and ‘Obsession’. Consensus sequences for each clone were assembled and annotated to identify intron regions by alignment to transcript sequences; coding regions were confirmed and annotated using InterProScan (Jones *et al*., 2014). Assembled sequences were identified as individual alleles, nucleotide and predicted protein sequences were aligned using the EMBL-EBI Clustal Omega multiple sequence alignment tool (Madeira *et al*., 2019), and alignments were visualized using Jalview 2.11.1.4 (Waterhouse *et al*., 2009).

Protein structures were predicted using RoseTTAFold (Baek *et al*., 2021), and the predicted secondary structure was added to the Clustal Omega allele sequence alignment using ESPript 3.0 (Robert & Gouet, 2014). Allele structures were used as a query search against the Protein Data Bank in DALI, and structure pairwise comparison was performed (Holm, 2020). Allele structures were further analyzed and aligned using UCSF Chimera (Pettersen *et al*., 2004). Allele expression was analyzed by mapping the variable coding regions to the RNA-seq data using Burrows-Wheeler Aligner software package (BWA-MEM) (Li & Durbin, 2009), and results were visually examined using the Integrated Genomics Viewer (Robinson *et al*., 2011).

## RESULTS

### Sequencing data reflect phylogenetic relatedness of MRI and SB22

We generated 12.8 million (MRI), 14.3 million (SB22), and 9.9 million (Sporangia) high quality paired-end reads per replicate for three controls (Table 1, Supporting Information Figure S1). Considering the increased complexity of infected samples, we doubled the sequence coverage and generated an average 24.8 million and 27.6 million high quality Illumina paired-end reads per replicate per infection sample for MRI and SB22, respectively (Table 1, Supporting Information Figure S1). All sequence data were deposited at NCBI under GEO NUMBER: GSE111387.

**Table 1.** Summary of assembled transcripts by timepoint.

| Dataset (organism) | Read pairs (millions) | Mapping Percentage | Average base quality | Number of Genes | Average gene length | Average coverage |
| --- | --- | --- | --- | --- | --- | --- |
| Sporangia (*P. belbahrii*) | 29.7 | 84.1 | 37.28 | 10144 | 1603 | 230.4 |
| MRI Water (*O. basilicum* MRI) | 42.7 | 62.0 | 37.20 | 66930 | 893 | 66.4 |
| MRI 12 hpi (*O. basilicum* MRI, *P. belbahrii*) | 75.9 | 63.3 | 37.21 | 81873 | 897 | 98.1 |
| MRI 24 hpi (*O. basilicum* MRI, *P. belbahrii*) | 75.3 | 63.3 | 37.22 | 89302 | 864 | 92.6 |
| MRI 48 hpi (*O. basilicum* MRI, *P. belbahrii*) | 77.0 | 63.1 | 37.23 | 85832 | 860 | 98.7 |
| MRI 72 hpi (*O. basilicum* MRI, *P. belbahrii*) | 69.8 | 63.4 | 37.22 | 78061 | 909 | 93.5 |
| SB22 Water (*O. basilicum* SB22) | 38.3 | 62.9 | 37.18 | 52844 | 977 | 69.9 |
| SB22 12 hpi (*O. basilicum* SB22, *P. belbahrii*) | 85.9 | 63.9 | 37.18 | 74918 | 934 | 117.6 |
| SB22 24 hpi (*O. basilicum* SB22, *P. belbahrii*) | 83.2 | 63.3 | 37.19 | 72699 | 949 | 114.5 |
| SB22 48 hpi (*O. basilicum* SB22, *P. belbahrii*) | 75.9 | 65.2 | 37.15 | 84478 | 870 | 100.9 |
| SB22 72 hpi (*O. basilicum* SB22, *P. belbahrii*) | 86.6 | 66.6 | 37.20 | 80675 | 903 | 117.7 |
| MRI Combined Assembly | 370.4 | 64.8 | 37.22 | 133,441 | 765 | 352.6 |
| SB22 Combined Assembly | 399.6 | 66.0 | 37.19 | 118,296 | 692 | 477.7 |

In total, 240.2 million paired-end reads were used to generate the MRI Combined Assembly, containing 341,633 unique transcripts corresponding to 133,441 genes called by Trinity. The SB22 Combined Assembly was generated using 263.9 million paired-end reads and contained 118,296 genes and a total of 322,696 unique transcripts. The MRI and SB22 plant-only control assemblies contained 66,930 and 52,844 genes respectively, and the sporangia control contained 10,144 assembled genes. As expected, more genes were assembled in infected samples, representing both host and pathogen transcripts, genes expressed only during infection, and assembly errors (fragmented sequences) introduced as transcriptome complexity increased.

Though fewer reads were sequenced for both water and sporangia control samples, the sporangia control produced the highest sequence coverage (>90x) and longest average gene length (1,603 bp) for assembled genes, likely due to the smaller genome/transcriptome size of the (inactive) pathogen. The average assembled gene length of *O. basilicum* transcripts from a previously published transcriptome was 1,363 bp (Rastogi *et al*., 2014), larger than our average assembly size. The average gene size of the oomycete *Phytophthora infestans* was 1,523 bp (Haas *et al*., 2009), roughly equivalent to the sporangia control assembly.

To assess the genetic diversity between MRI and SB22, we performed a BLAST search between MRI and SB22 water control assemblies and identified 20,943 reciprocal hits, likely representing orthologs between these two plants. These orthologs are highly similar with an average pairwise sequence identity of 98.66% (Figure 1B). We further assessed the genetic similarity between the cultivars by conducting a phylogenetic analysis of nine protein-coding chloroplast genome orthologs across members of the asterid clade, to which *O. basilicum* belongs, and with *A. thaliana* and *S. oleracea* set as outgroups (Rastogi *et al*., 2015). The analysis showed high sequence conservation among the *Ocimum* spp. with MRI and SB22 grouped together (Figure 1C). We anticipate that unique genes or differentially regulated genes are likely to contribute to the MRI and SB22 phenotypic variations.

Using expression profiles in the individual assemblies, MRI transcripts can be divided into 36,414 predicted plant transcripts (present in the MRI water control), 9,988 predicted pathogen transcripts (present in the sporangia control), and 29,502 infection unique transcripts (absent in both plant and pathogen controls) (Table 2). Similarly, the SB22 transcripts include 31,702 transcripts with a plant origin, 9,426 transcripts of pathogen origin, and 26,486 transcripts uniquely present in the infection samples. Consistent with SB22 susceptibility, we saw a significant increase in the number of expressed pathogen genes in the susceptible host, observing an almost 20-fold increase from 12 to 72 hpi compared to only a two-fold increase for MRI (Figure 2). Measuring total pathogen mRNA abundance (the number of reads mapping to roughly 4,553 pathogen genes), we detected a 43-fold increase in mapped pathogen reads from the SB22 72 hpi samples relative to the MRI 72 hpi samples (Figure S2).

**Figure 2.**
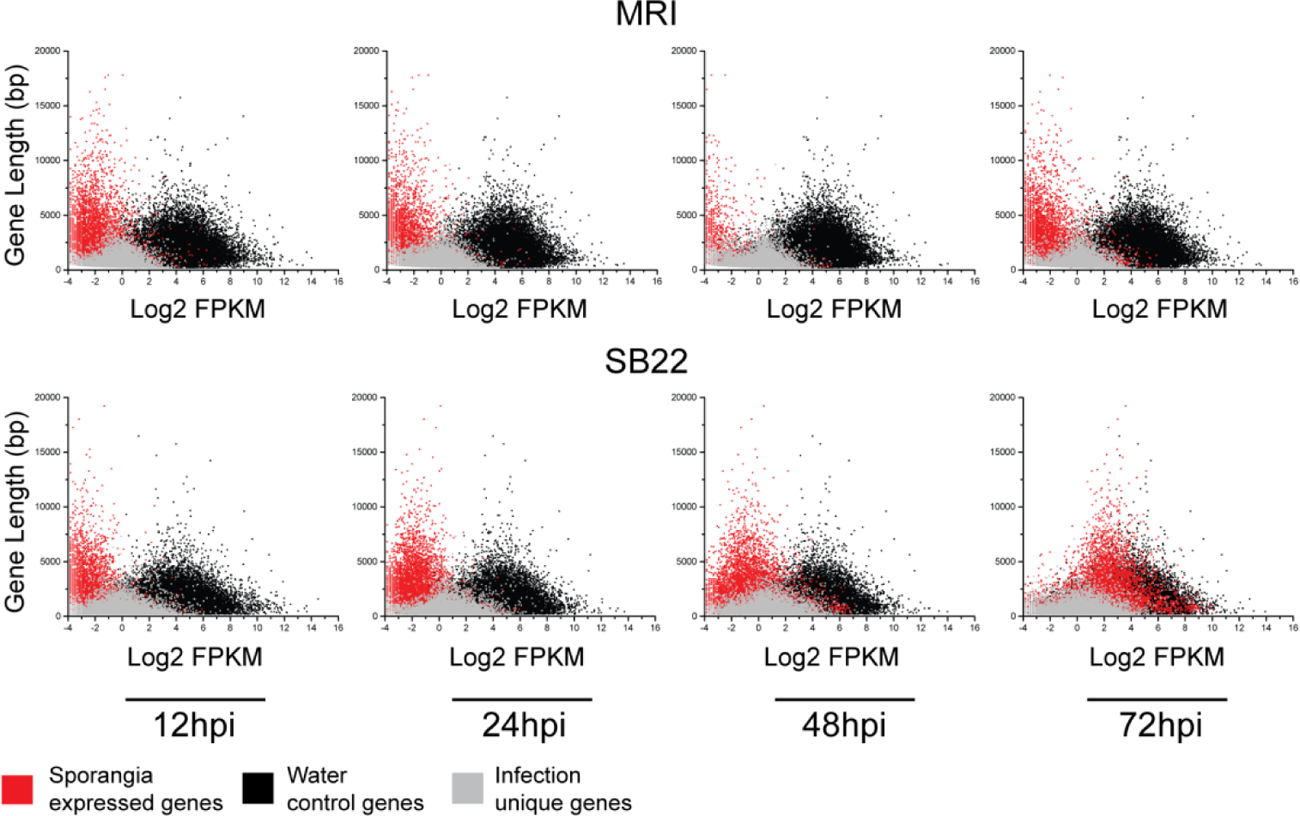
Plant and pathogen gene expression profiles from clustering of the MRI Combined Assembly. Clustering was done using all datasets mapped to the MRI combined assembly. Cluster numbers are displayed above each cluster image. Dataset and replicate are listed on the bottom x-axis. Sporangia expressed gene expression is in red, infection unique gene expression is in gray and gene expression in the mock infected water controls are in black for both MRI and SB22.

**Table 2:** Expressed genes across timepoints

| Dataset | Number of Genes FPKM > 1 |  |  |  |
| --- | --- | --- | --- | --- |
|  | Total Genes | Plant Control | Sporangia Control | Infection Unique |
| MRI Water | 36,528 | 36,414* | 0 | 0 |
| MRI 12 hours | 39,349 | 29,247 | 472 | 9,534 |
| MRI 24 hours | 40,171 | 29,074 | 389 | 10,606 |
| MRI 48 hours | 40,973 | 28,852 | 179 | 11,840 |
| MRI 72 hours | 39,279 | 28,365 | 825 | 9,987 |
| Sporangia | 10,102 | 0 | 9,988* | 0 |
| **Total unique** | **76,018** | **36,414** | **9,988<sup>+</sup>** | **29,502** |
| SB22 Water | 31,794 | 31,702* | 0 | 0 |
| SB22 12 hours | 35,561 | 25,770 | 274 | 9,431 |
| SB22 24 hours | 36,732 | 25,584 | 699 | 10,364 |
| SB22 48 hours | 40,216 | 25,746 | 1,788 | 12,598 |
| SB22 72 hours | 40,838 | 25,414 | 5,375 | 9,962 |
| Sporangia | 9,518 | 0 | 9,426* | 0 |
| **Total unique** | **67,706** | **31,702** | **9,426<sup>+</sup>** | **26,486** |

\*- When the data was split into 3 sets 114 MRI points and 92 SB22 points were removed due to ambiguity
+ - The numbers displayed in in the chart above are total genes in each category as split by FPKM based separation after mapping to each master assembly. If the gene had a non-zero FPKM in the sporangia samples it was included in the "sporangia" sample. The difference in gene count is a reflection of mapping to two seperate assemblies.

### Clustering analysis highlights transcripts with potential functions involved in host - pathogen interactions

Based on overall ortholog identity, we were confident that SB22 reads from orthologous genes would align to the MRI assembly. Using the MRI Combined Assembly as a reference, we mapped all the SB22 and MRI reads using RSEM with default parameters. Initial coarse clustering using all genes from MRI and SB22 mapped data resulted in 12 clusters with on average 2,810 genes per cluster (Supporting Information Table S2). Three clusters that lacked consistency among biological replicates were not included in further analyses. Three clusters (A1, A2 and A3) characterized transcripts primarily belonging to the pathogen (Figure 3 panel A), as all transcripts showed significant expression in sporangia pathogen control samples (Figure 3 grey bar in the middle), but no expression in both water-inoculated plant control samples (Figure 3 two blue bars representing MRI-water only and SB22-water only).

**Figure 3.**
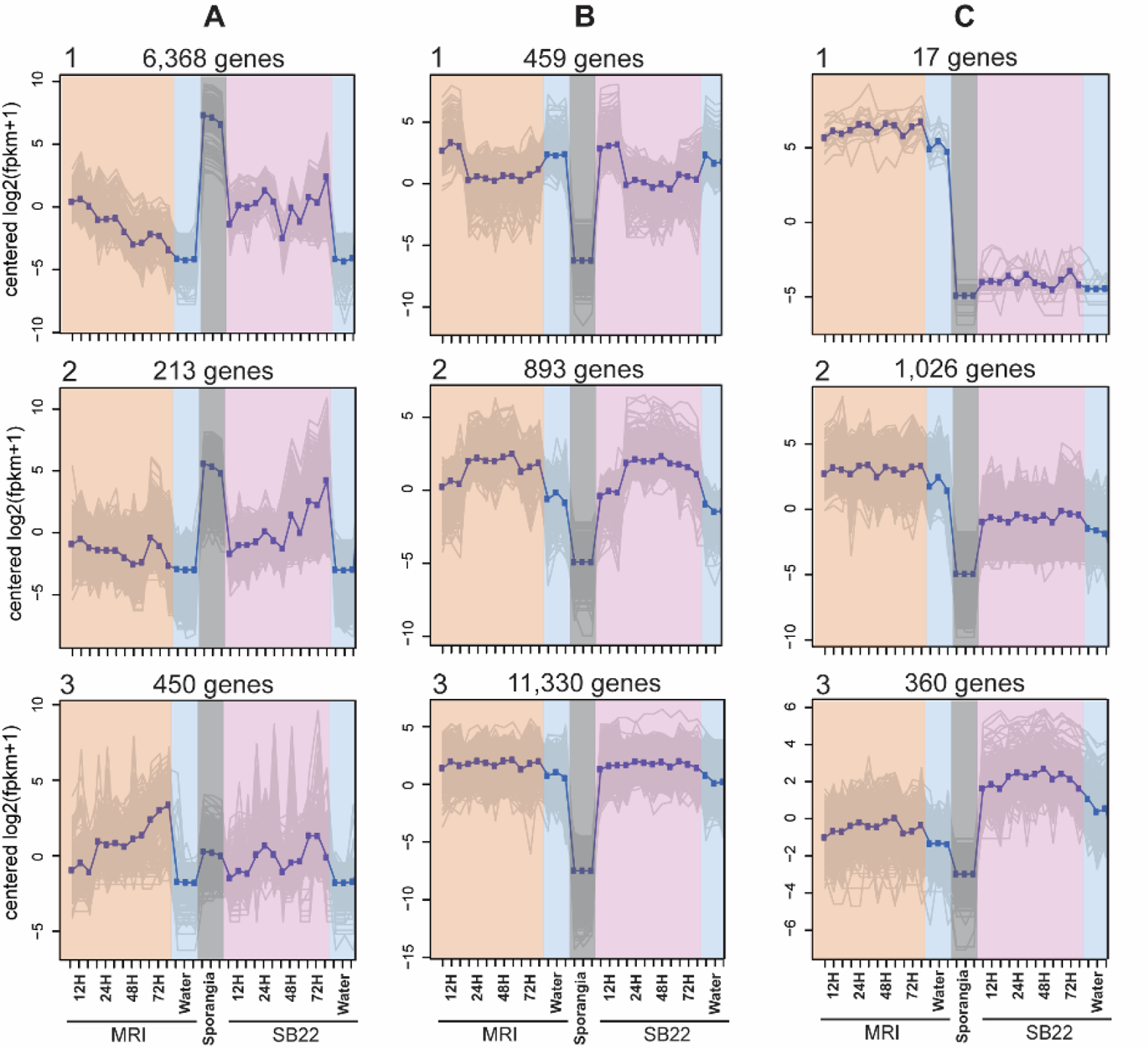
Clusters with expressed plant genes. Nine gene clusters produced by coarse clustering are displayed here. Expression value, y-axis, is on a log2 scale. Three replicates for each dataset are represented by tick marks on the bottom x-axis, with mean data plotted as dark blue points and gray shading representing the standard deviation. Datasets from MRI are highlighted in orange with 3 replicates from 12 to 72 hpi, SB22 datasets are colored pink with 3 replicates from 12 to 72 hpi, both water controls are labeled blue (MRI left, SB22 right), and the pathogen sporangia control data is colored gray. Column A, pathogen gene clusters as all transcripts showed significant expression in sporangia pathogen control samples but were absent from plant and infected plant samples. Columns B and C gene clusters contain primarily plant transcripts as all transcripts showed significant expression in plant and infected plant samples but were absent from sporangia pathogen control samples. Column B, plant genes expressed similarly in both cultivars. Column C, plant genes with different profiles between cultivars.

Six clusters containing primarily plant transcripts were identified. All transcripts showing significant expression in plant and infected plant samples that were absent from sporangia pathogen control samples were filtered (Panels B and C). The remaining genes from the six plant gene clusters represent plant transcripts during pathogen challenge. Transcripts within three clusters, B-1, B-2 and B-3, had comparable expression profiles between MRI and SB22, indicating conserved functions between the two different plant hosts. Both B-1 and B-2 clusters show a pattern consistent with a 12-hour shift in photoperiod, but these clusters respond in opposite directions. Cluster B-1 was upregulated at 24, 48, and 72 hpi and was enriched for metabolism, oxidation-reduction, and photosynthesis functions. No GO terms were significantly enriched in cluster B-2 which showed downregulation at 24, 48, and 72 hpi; however, of those GO terms annotated by Blast2Go metal ion functions were predominant. The largest cluster, B-3, roughly represents stably expressed plant genes. Cluster B-3 is enriched for many categories including various metabolic processes, protein modification, and protein localization, among others.

Transcripts in three clusters, C-1, C-2, and C-3 displayed differential regulation responses between MRI and SB22 during pathogen challenge (Figure 3B). In Cluster C-1, MRI expression is high but almost completely absent from the SB22 transcriptome. In Cluster C-2, MRI expression was higher than SB22 and was enriched for genes related to defense response, response to stress, response to stimulus, and DNA integration. In Cluster C-3, the expression of MRI genes is instead lower than SB22. No GO terms were enriched in cluster C-3.

### MRI unique expressed genes include NLR, RLK and secondary metabolic enzymes

To understand potential mechanisms underlying the resistance, we repeated the clustering with higher stringency (p=20, see methods for details) resulting in 188 clusters. Eight clusters were chosen as they were expressed in MRI, minimally expressed in SB22, and showed no expression in the sporangia control. A comprehensive filtering process of the eight clusters resulted in a total 369 MRI unique candidate genes. These MRI unique candidate genes can be grouped into secondary metabolic enzymes (22 genes), immunity related genes including 22 nucleotide-binding site leucine-rich repeat (NLR) genes and 25 receptor-like kinases (RLK) or receptor-like proteins and others.

Detecting secondary metabolic enzymes as MRI unique genes is expected as these two basil plants produce distinct secondary compounds. For instance, SB22 accumulates a significant amount of eugenol, while MRI predominantly accumulates methylchavicol (Rob Pyne, unpublished data). This distinct chemotype prevents MRI from immediate commercial use. Examining twenty-five secondary metabolite related genes predicted as MRI unique genes, we found enzymes related to secondary metabolites which specifically differentiate the MRI and SB22 chemotypes, including cinnamate p-coumarate carboxyl methyltransferase, enzymes involved in anthocyanin biosynthesis, and chavicol/eugenol O-methyltransferase, the enzyme that catalyzes the conversion of chavicol to methylchavicol, as would be predicted from the chemotypes.

Unique expression of RLKs and NLRs, both immunity related protein families, in MRI is of particular interest. Basic plant immunity consists of pattern-triggered immunity (PTI) and effector-triggered immunity (ETI) (Jones & Dangl, 2006; Cui *et al*., 2015). Plant PTI uses receptor-like proteins/kinases (RLPs/RLKs), a large gene family (Chern *et al*., 2016; Mendy *et al*., 2017) that have roles as sensors of microbe-associated molecular patterns (MAMPs) and induce downstream defense reactions (Jones & Dangl, 2006; Dodds & Rathjen, 2010; Bigeard *et al*., 2015; Zhou & Zhang, 2020). Plant ETI employs an intracellular nucleotide-binding site and leucine-rich repeat domain receptors (NLRs) that play roles in sensing effector proteins secreted by pathogens and regulating downstream defense signaling (Jones & Dangl, 2006; Cui *et al*., 2015; Cesari, 2018; Monteiro & Nishimura, 2018; Wang & Chai, 2020).

To investigate basil PTI, we further characterized the RLK BLAST hits. The best RLK candidate is transcript mri_comp170662, which encodes a full-length malectin-like RLK. Some predicted receptor-like kinases (RLK) as truncated fragments, such as three transcripts containing only the leucine-rich repeat (LRR) domain, two containing only protein kinase domains, and two that were too short to have domain affiliation. Two transcripts contained an RLK-like domain involved in antifungal and salt tolerance (Sawano *et al*., 2007; Zhang *et al*., 2009). Using the full-length top candidate RLK transcript mri_comp170662, we identified 12 and 10 homologs in the MRI and SB22 assemblies, respectively. Ten SB22/MRI orthologous pairs can be readily identified between these two sister cultivars based on a phylogeny (three pair members lacking a malectin domain were excluded from the tree) (Figure 4a). Three MRI orphan transcripts including this top RLK candidate mri_comp170662 showed increased expression at 12 and 24 hpi compared to the water control (Figure 4b). This RLK candidate contains LRR, Malectin and kinase domains (Figure 4c).

**Figure 4.**
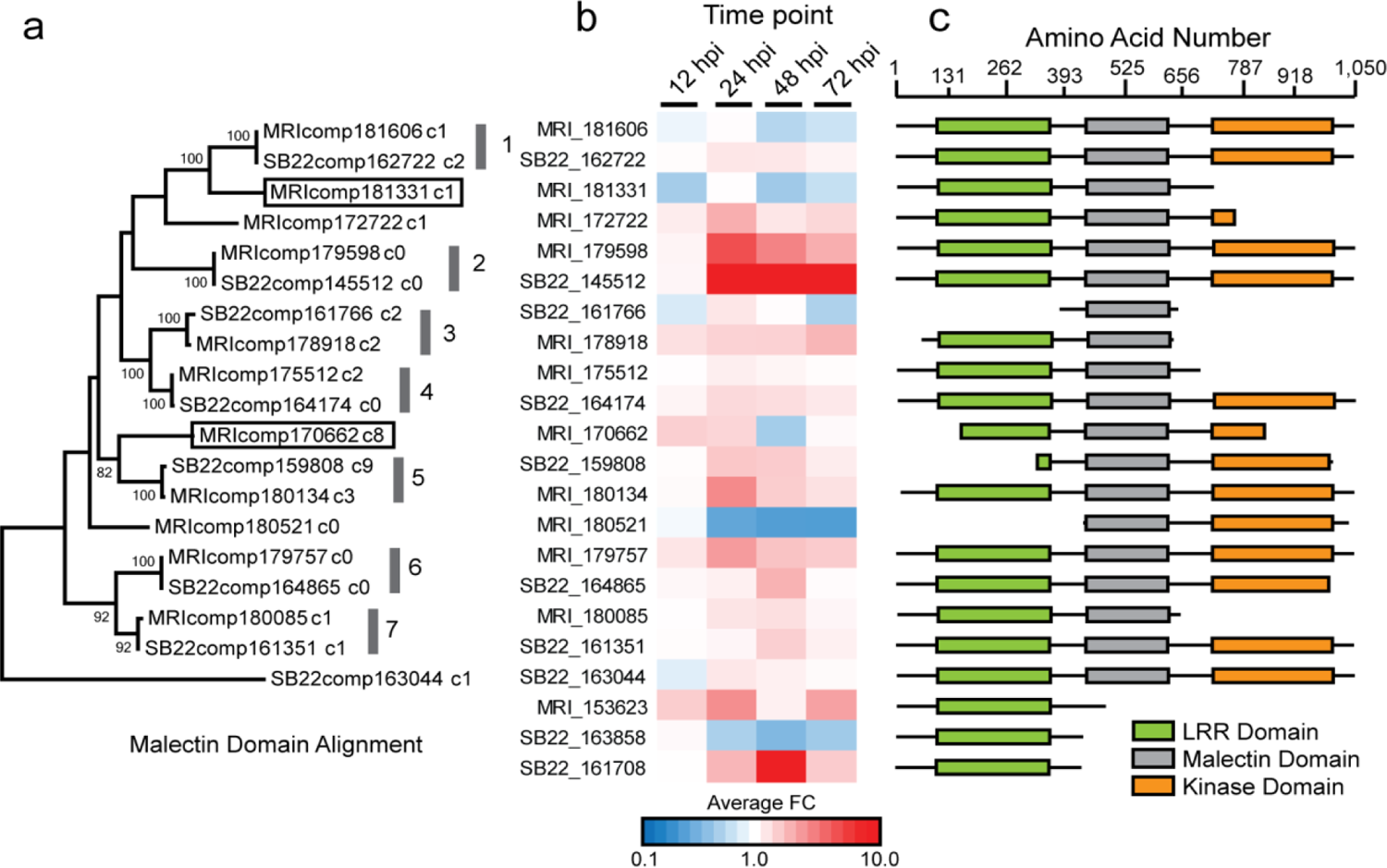
Malectin-like RLK proteins in MRI and SB22. (a) Alignment of 19 of 22 RLK proteins by their conserved malectin domain. Gray bars with numbers represent orthologous pairs and boxes indicate sequences absent from the SB22 cultivar. (b) Fold change compared to water for MRI and SB22 RLKs across infected plant samples. (c) Protein domain structure of 22 RLK hits generated from translated nucleotide sequences. Transcript IDs are those of the adjacent fold change row.

### NLRs upregulation in MRI during infection

To identify unique MRI NLR resistance genes involved in ETI, we focused on Cluster C-1 where all 17 transcripts are highly expressed in MRI upon infection but almost absent from the SB22 transcriptome. Two transcripts, comp_178221_c0 and comp_160460_c0, are putative NLR resistance genes encoding a late blight resistance protein homolog R1A (gi:848916018 and gi:848932751) from spotted monkey flower (*Erythranthe guttata*), which belongs to the order Lamiales including basil.

Members of the plant NLR protein family have been characterized as sensors, recognizing specific microbial effectors in response to ongoing host-pathogen coevolution, or as helpers involved in signal transduction (Wu *et al*., 2017). These proteins are highly conserved in eukaryotic immune responses (Wu *et al*., 2017). NLRs contain a central nucleotide-binding domain and a C-terminal leucine-rich repeat region that confers specificity to the receptor (Wu *et al*., 2017; Prigozhin & Krasileva, 2020). There are three subfamilies of NLRs defined by presence of one of three functional N-terminal domains: Resistance To Powdery Mildew 8 (RPW8), Coiled-Coil (CC), or Toll/Interleukin-1 Receptor homology (TIR) (Prigozhin & Krasileva).

The coiled-coil domain NLR subfamily (CC-NLRs) comprises several well-characterized intracellular receptors, including Recognition of Peronospora Parasitica 1 (RPP1) genes conferring resistance to downy mildew in *A. thaliana* (Krasileva *et al*., 2010). CC-NLRs are characterized by an N-terminal CC domain, which has been associated with oligomerization in characterized CC-NLRs including barley mildew locus A MLA10 and *A. thaliana* ZAR1, an NLR that polymerizes to form a plant “resistosome” during the immune response (Maekawa *et al*., 2011; Adachi *et al*., 2019).

We focused on validating our computational prediction and further examined our top candidate NLR transcript comp160460_c0 identified in Cluster C-1. Transcript comp_160460_c0, MRI Resistance gene 1 (MRI-R1), encodes a full-length CC-NLR protein of 887 aa with all three functional domains. This NLR candidate is uniquely expressed and differentially upregulated in MRI in the presence of the pathogen (Figure 5A). Based on mapping results and expression analyses, while MRI-R1 is expressed in the control samples, its expression is significantly increased in the pathogen-inoculated samples throughout the time-course of infection. Specifically, MRI-R1 was 2-fold upregulated between 12 and 24 hpi and upregulated expression was maintained 48- and 72-hpi. Mapping the SB22 infected assemblies against the MRI-R1 transcript sequence as a reference confirmed that there was no detectable expression of MRI-R1 detected in susceptible samples.

**Figure 5.**
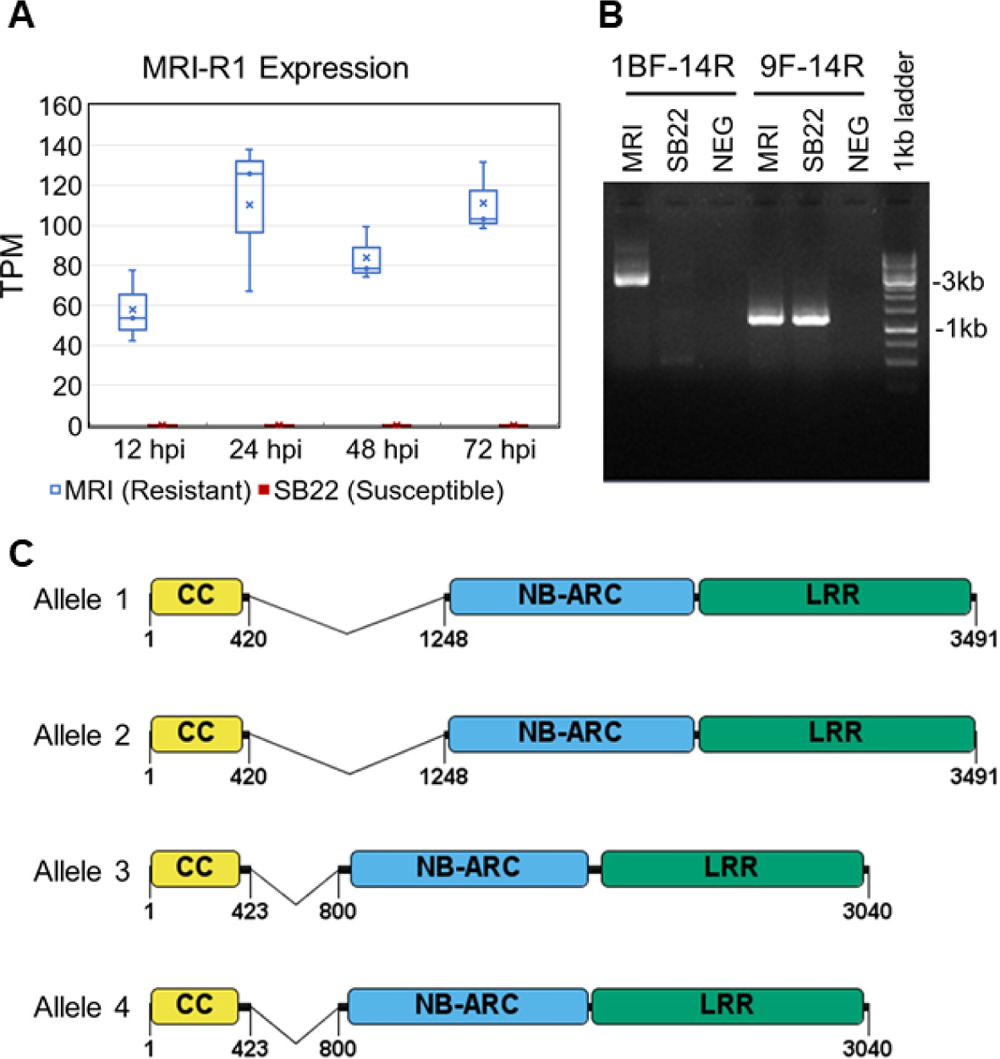
MRI-R1 unique presence, expression, and alleles in MRI. A) Expression of MRI-R1 in *P. belbahrii*-inoculated MRI and SB22 cultivars expressed in TPM (transcripts per million). B) Unique amplification of MRI-R1 from MRI (3673bp) using the external 1BF forward and 14R reverse primers. Internal primer, 9F, paired with 14R produces amplicons of expected size 1206bp in both MRI and SB22. The negative control (NEG) is water in place of gDNA template. C) Gene models of the 4 MRI-R1 alleles with domains and subdomains predicted by InterProScan colored as: coiled coil (gold), NB-ARC (blue), and leucine-rich repeat (green). Introns between the coiled coil and NB-ARC domains are represented as a single line.

To determine whether MRI-R1 activity could be attributed to presence/absence polymorphism between MRI and SB22 or the differential regulation at transcriptional level, we designed primers to amplify the coding region of the full-length MRI-R1 transcript using PCR. Forward primer 1BF flanks the 5’ end of the coding sequence, and reverse primer 14R targets the 3’ end of the gene. This primer pair produced an amplicon of the MRI-R1 gene from MR1, but not from SB22 or the water template negative control (Figure 4B). Internal primers were designed to further examine MRI-R1 like genes in MRI and SB22 (Supporting Information Table S1). PCR using internal forward primer 9F, which flanks the NB-ARC domain, paired with 3’ end primer 14R, resulted in gene amplification from MRI and potentially off-target or ortholog amplification from SB22 (Figure 5B). These results suggest that the full MRI-R1 gene is unique to MRI, but that there is some partial sequence conservation in a similar gene of unknown functional status in SB22. Thus, MRI-R1 represents another case of R gene polymorphism among closely related organisms.

The MRI-R1 amplicon detected in the MRI gDNA appeared to be approximately 800bp larger than predicted from the RNA transcripts, indicating the potential presence of a non-coding intronic region (Figure 5B). We also observed sequence polymorphisms among MRI-R1 transcripts with 6 different isoforms predicted for the same gene (2 with full coding sequences, comp160460_c0_seq2 and comp160460_c0_seq5). To investigate transcript polymorphisms we isolated and sequenced individual clones from the PCR products. A total of 12 full-length amplicons were cloned from three MRI plants. Due to the length of the transcript and to help validate the sequences, between 12 to 25 overlapping sequences were generated. Assembly and annotation of these MRI-R1 sequences revealed 4 separate alleles of the gene, supporting the allelic polymorphisms observed in the transcriptomic data (Figure 4C). The pairwise nucleotide sequence identity among the four MRI-R1 alleles range from 80% to 96.45%. Alleles 1 and 2 share an identical 827 nucleotide intron sequence while alleles 3 and 4 share an identical 376 nucleotide intron sequences, which accounts for the size discrepancy in the agarose gel electrophoresis result (Figure 5B).

Analysis of the protein sequences using InterProScan (Jones *et al*., 2014) showed that all four alleles contain the CC, NB-ARC, and LRR domains with canonical functional motifs. The N-terminus of all four alleles begins with an identical MADA motif, a functional motif conserved in approximately 20% of CC-NLR immune receptors across distantly related plant species (Bentham *et al*., 2018; Adachi *et al*., 2019). The MADA motif has been shown to be necessary for *Nicotiana benthamiana* NRC4 cell death like the ZAR1 resistosome (Adachi *et al*., 2019). Similarly, the EDVID motif known to be involved in self-association, direct interactions with cofactors and, in some cases, cell death signaling resulting in a hypersensitive response (Bentham *et al*., 2018), is also present in all 4 alleles. The NB-ARC domain of all four alleles also contains the Walker A (P-loop) motif (GMFGLGKT) (Ramakrishnan *et al*., 2002), which is critical for nucleotide binding (Steele *et al*., 2019). Also present in the four alleles is the MHD-type motif IHD, which has been shown to be involved in inhibition of autoactivation of R proteins in the absence of a pathogen (van Ooijen *et al*., 2008). The MHD motif is proposed to act as a molecular switch for R protein activation, and the histidine and aspartate residues are the most highly conserved across R proteins, with the histidine occupying a critical position in an ADP-binding pocket (van Ooijen *et al*., 2008). The most variable regions among these 4 alleles are in and between the CC and NB-ARC domains (Figure S3).

Protein structural models of all four alleles were generated in the Robetta server utilizing RoseTTAFold, a top-ranked deep-learning based protein structure prediction method (Baek *et al*., 2021; Du *et al*., 2021). The resulting allele protein structures were queried using a distance matrix alignment in the DALI server to search for the closest structural homologs (Holm, 2020).

The resulting top hits for MRI-R1 included plant proteins such as NB-ARC domain from the tomato immune receptor NRC1 (6S2P), LRR receptor-like serine/threonine-protein kinase FLS2 (4MN8), and LRR receptor-like serine/threonine-protein kinase GSO1 (6S6Q). The top structural homologs also included human and animal LRR proteins such as Leucine-rich repeat transmembrane neuronal protein 2 (5Z8X), Dimeric bovine tissue-extracted decorin (1XCD), and human osteomodulin (5YQ5). The top ten structural homologs ranged in Z scores of 18-22.2 (with Z scores above 20 indicating definite homology, and above 8 indicating probable homology), and average deviation in distance between aligned Cα atoms in 3-D superimpositions, indicated by root mean-squared deviation (RMSD) ranged from 2.2-8.5 angstroms (Supporting Information Table S3) (Holm, 2020).

Structural modeling and homology comparison of each allele were used to refine the initial sequence-based boundary predictions of CC, NB-ARC, and LRR domains (Figure 6A). All 4 allele structural models were aligned, revealing conservation of predicted functional domain structures and active sites. As expected, based on high sequence identity, all four alleles shared high structural similarity, with slight differences in domain boundaries predicted by the models (Figure 5B). Allele expression analysis showed a clear bias for expression of MRI-R1 Allele 1 by MRI in response to basil downy mildew infection.

**Figure 6.**
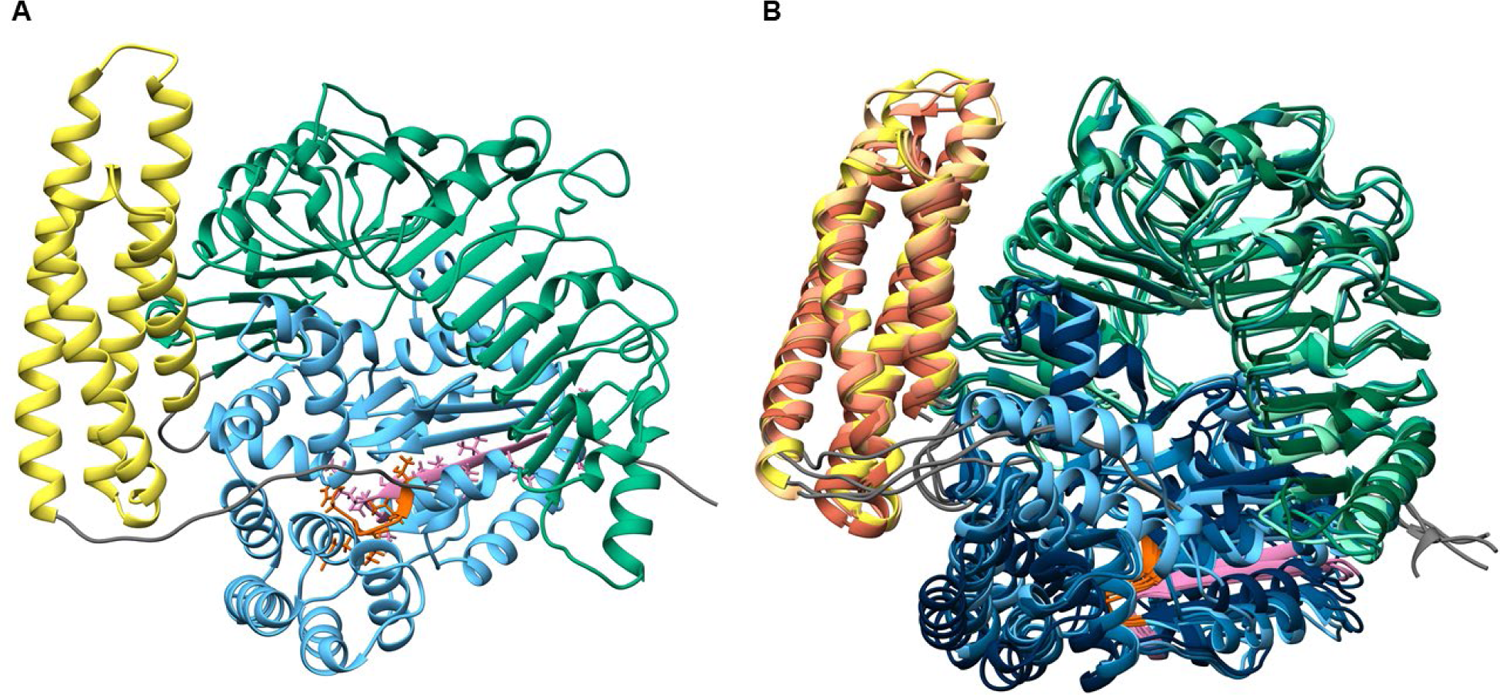
MRI-R1 protein structural modeling and alignment. a) Predicted protein structure of MRI-R1 allele 1 with domains and motifs colored from N to C terminus as follows: coiled coil domain in yellow, NB-ARC domain in blue, Walker A motif in red, Walker B motif in purple, and LRR domain in green. b) Alignment of predicted structures for all 4 MRI-R1 alleles.

### The presence of MRI-R1 in two Backcross progenies ‘Devotion’ and ‘Obsession’

To assess the contribution of MRI-R1 in the four new downy mildew resistant (DMR) basil cultivars that integrated MRI resistance genes, we tested each for the presence of the MRI-R1 gene. A partial sequence of ObActin, the ubiquitous positive control, was successfully amplified from both BDM-susceptible SB22, BDM-resistant MRI, and all four DMR cultivars, while no amplification was detected from the negative water template control (Figure 7). Interestingly, MRI-R1 was detected in only two, ‘Devotion’ and ‘Obsession’, out of the four new DMR cultivars using primers designed to amplify the full coding region of the gene (Figure 7). The two amplicons are estimated to be 3064 bp and 3515 bp, corresponding to allele 4 and allele 1 for ‘Devotion’ and ‘Obsession’, respectively. These amplicon sizes include 24bp of the 5’ UTR (beginning with primer 1BF), and the 827bp intron (alleles 1 and 2) or the 376 bp intron (alleles 3 and 4). The six individual clones selected and sequenced from ‘Devotion’ and ‘Obsession’ were uniform, indicating that only one allele of MRI-R1 was passed from MRI to these offspring.

**Figure 7.**
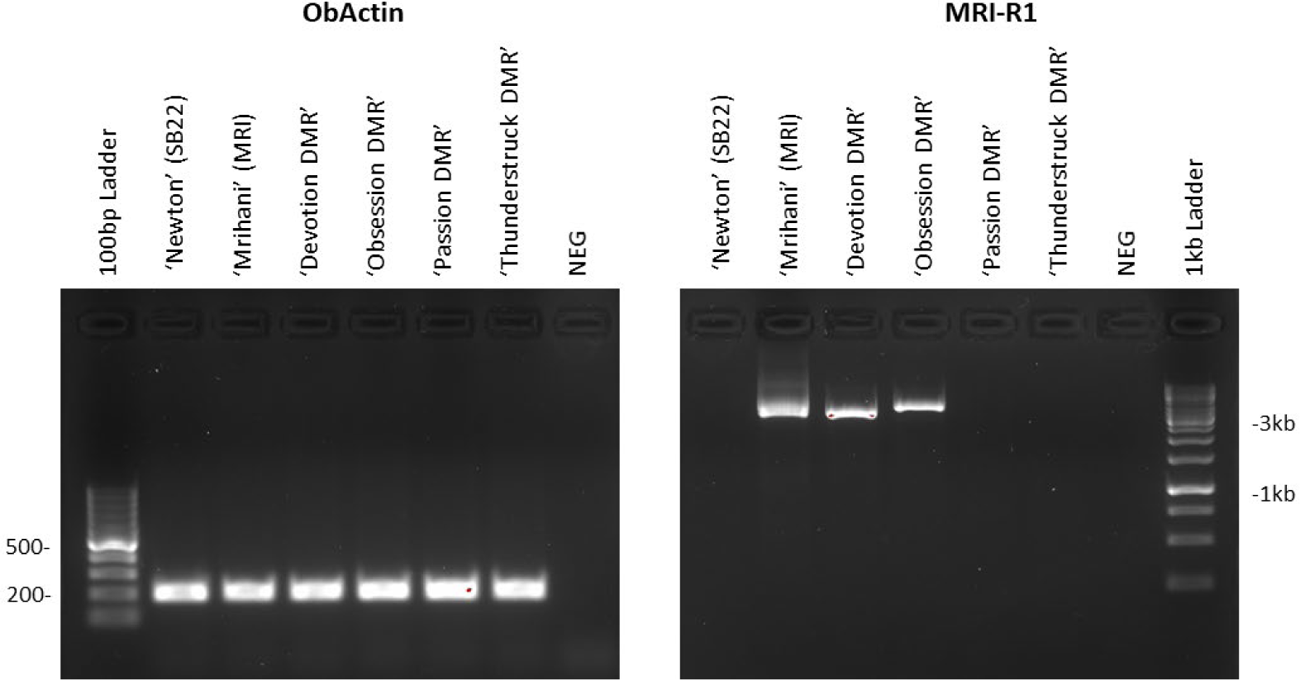
MRI-R1 detection in DMR offspring. Gel electrophoresis showing amplification of ObActin fragment positive control from all basil cultivars tested on the left, MRI-R1 amplification from ‘Mrihani’ (MRI), ‘Devotion’, and ‘Obsession’ on the right using the external 1BF forward and 14R reverse primers. The negative control (‘control’) is water in place of gDNA template.

**Figure 8.**
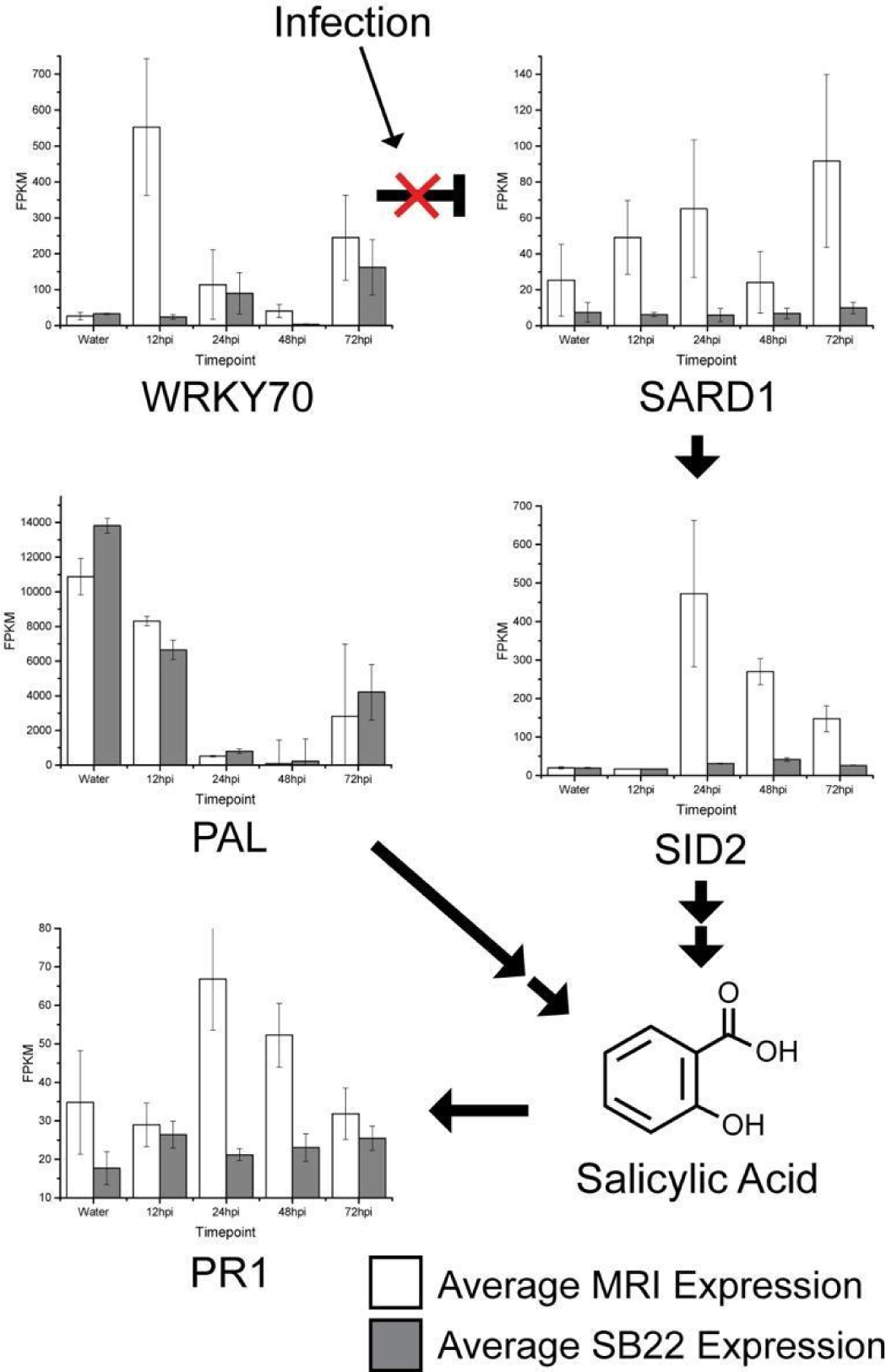
MRI and SB22 diverge transcriptionally at salicylic acid synthesis. Bar graphs are the average and standard deviation of three replicates. White bars indicate MRI data and gray bars indicate SB22 for each timepoint. Infection releases the repression WRKY70 has on SARD1 expression. Double arrows leading from SID2 and PAL indicate more than one step to the SA molecule. The most striking difference was observed among genes required for the synthesis of salicylic acid (Figure 8). Plants possess two biosynthesis pathways to synthesize SA, both starting from chorismate, but subsequent steps involve either isochorismate synthase (ICS) or SALICYLIC ACID INDUCTION DEFICIENT 2 (SID2) (Wildermuth *et al*., 2001) and phenylalanine ammonia-lyase (PAL) (Olsen *et al*., 2008). There is no significant difference in PAL expression between the two cultivars, but we did observe a drastic induction of ICS 24 hpi in MRI. For the water control and 12 hpi samples, there was no significant difference in ICS expression between MRI and SB22. Compared to SB22, ICS expression in the MRI cultivar is 15, 6 and 5-fold higher at 24, 48 and 72 hpi, respectively.

These four new DMR Cultivars were selected from genetic breeding efforts. Briefly, F1 progeny were generated through the MRI (female)×SB22 (male) cross, exhibiting dominant gene action (Pyne *et al*., 2015). An F2 family was generated after F1 self-pollination and a single resistant individual RUMS469-11 was selected for hybridization with elite sweet basil inbred line ‘SB13’, which demonstrates downy mildew and Fusarium wilt tolerance. Twenty individuals achieving the highest category of reduced disease severity from the RUMS469-11 (female)×SB13 (male) cross were self-pollinated to generate full sibling families evaluated for response to downy mildew. The DMR ‘Devotion’, ‘Obsession’, ‘Passion’ and ‘Thunderstruck’ were selected from these inbred lines. We anticipate these four new selected DMR cultivars should inherit genetic resistance genes from both MRI and SB13.

### Differential upregulation of salicylic acid biosynthesis pathways in MRI

Both NLR and RLK genes interact with hormone signaling pathways to activate host defense (McHale *et al*., 2006). To understand the involvement of plant hormone signaling pathways involved in susceptibility and resistance responses, we examined the expression of genes involved in ethylene, jasmonic acid (JA), abscisic acid (ABA), jndole-3-acetic acid (Auxin), gibberellic acid (GA), and salicylic acid (SA) based on *A. thaliana* annotation (1e-20, sequence similarity >40%). No significant differences were observed between the MRI and SB22 pattern of expression in ethylene, jasmonic acid, abscisic acid, or gibberellic acid pathway genes (Figure S4, Table S2). We saw a difference in expression profile for two of five auxin genes; YUC1 and TAA1 both were upregulated at early timepoints in MRI, however they were not statistically significantly differentially expressed (p value > 0.05).

SARD1 is upstream of SID2 in the SA synthesis pathway, and the average SARD1 expression across all time points is 7-fold higher in MRI compared to SB22 and has roughly 2-, 2.5-, 0- and 3.6-fold increases at 12, 24, 48 and 72 hpi compared to the water control. Similarly, PR1, a commonly used marker gene downstream of SA synthesis, is significantly induced in MRI upon pathogen challenge starting at 24 hpi, confirming the unique increase of SA in MRI upon pathogen challenge.

Together, our data suggest that MRI-derived BDM resistance involves SA. The most significant change occurred 24 hpi, one day after the encounter of the plant with the pathogen. WRKY70 is known to repress SARD1 expression in the absence of pathogens and is required for the activation of some defense genes (Li *et al*., 2004; Zhou *et al*., 2018). In MRI, we saw a strong induction in the transcription of WRKY70 at 12 hpi, increasing expression of SARD1 from water to 24 hpi, and a 23-fold rise in the relative normalized expression of SID2 at 24 hpi relative to 12 hpi. In SB22, we observed only a slight increase in WRKY70 expression which was delayed until 24 hpi, and SID2 expression rose only 2.5-fold by 24 hpi and remained effectively stable at later time points. Similarly, we saw a roughly 2-fold induction in the expression of PR1 in MRI between 12 hpi and 24 hpi, consistent with the upregulation of SID2, while at the same time there was no increase in PR1 expression in SB22 between 12 and 72 hpi.

## DISCUSSION

Here we report the results of transcriptomic sequencing of basil cultivars infected with *Peronospora belbahrii* with the goals of identifying genes conferring resistance and understanding what pathways are involved in susceptibility. We sequenced 33 datasets to a reasonable coverage, likely capturing most transcripts, although total gene length was shorter than a previous study in sweet basil (Rastogi *et al*., 2014), and Trinity’s estimate of gene count in the plant samples was greater than expected. Nonetheless, we identified roughly 21,000 orthologous genes between cultivars with overall high sequence conservation consistent with closely related individuals. Phylogenetic analysis supports the hypothesis that SB22 and MRI are closely related, interspecific genotypes, which was previously shown using SSRs to confirm conservation of the orthologs in the two breeding parents (Pyne *et al*., 2018). These results underscore the close relationship between cultivars and their sexual compatibility, likely providing us with a pool of true genes shared in MRI and SB22. At the time of this study, no sweet basil genome was available for use in a reference-based assembly, though a draft genome for Genovese-type cultivar ‘Perrie’ has since been published (Gonda *et al*., 2020). Furthermore, several unique MRI gene candidate sequences were used to search the *O. basilicum* draft genome and no significant BLAST hits were retrieved.

Gene expression clustering identified patterns consistent with MRI unique infection-expressed genes. Annotated MRI unique candidates were found to be enriched for NLR, RLK, and secondary metabolic proteins. Analysis of the NLR and malectin-like RLK genes in both MRI and SB22 identified orthologous pairs and cultivar-unique genes. Members of the NLR and RLK families are known to act alongside or upstream of disease signaling pathways and serve as good candidate resistance genes. Secondary metabolite genes identified in the two cultivars correlate well with known chemotypic characteristics differentiating MRI and SB22. The top candidate NLR prediction is supported by PCR screening revealing its presence in resistant MRI and absence in susceptible SB22. This result was further strengthened with the cloning and sequencing-based confirmation of MRI-R1 alleles, as well as predicted amino acid sequence analysis and protein structure modeling supporting the hypothesis that this is a unique NLR likely involved in immune responses to pathogen infection.

The presence of MRI-R1 was detected in only two out of the four resistant cultivars, ‘Devotion’ and ‘Obsession’. However, recent quantitative trait loci (QTL) analysis detected at least two major genomic regions (LOD>4.0) that control DM resistance in the MRI x SB22 F2 mapping population (Pyne *et al*., 2017), suggesting that the predicted involvement of MRI-R1 in quantitative disease resistance may be redundant or have shared function(s) with other NLRs and/or RLKs. We cannot conclusively determine the functionality of the alleles simply based on the bias for MRI-R1 allele expression in MRI. The presence of Allele 1 in ‘Obsession’ and Allele 4 in ‘Devotion’ indicates that there may be functional redundancy, and further understanding of the conserved motifs in these alleles suggests that there may be interacting partners that have an impact on the activity of these proteins. Nevertheless, we are confident that our transcriptomic pipeline is powerful in detecting resistant genes involved in the host-pathogen interactions.

In addition to prediction of specific genes likely conferring resistance, this comparative transcriptomic approach was also valuable in revealing physiological mechanisms involved in basil downy mildew resistance. Salicylic acid signaling is an integral part of plant defense responses and has been demonstrated to be involved in defense against downy mildews and other biotrophic pathogens (Delaney Terrence P. *et al*., 1994; Mohr *et al*., 2010). Pathogen virulence strategies have developed to overcome and inhibit salicylic acid defenses, thus enhancing susceptibility to biotrophic pathogens such as downy mildew organisms (Caillaud *et al*., 2013, 2016). This suggests a likely role of salicylic acid signaling in MRI BDM resistance, though the specific mechanisms of signaling induction remain unknown. We hypothesize that multiple NLRs and RLKs are active and have interacting and/or redundant roles in mediating the SA signaling pathway in MRI. If this hypothesis is correct, utilizing multiple targets in a marker-assisted selective breeding program will be more effective and robust to pass resistance from parents to progeny in order to slow the evolution of new pathogen races.

This study has utilized mRNA sequencing over an infection time course to provide strong evidence linking susceptibility to known mechanisms which control defense responses, and prediction of genes regulating the resistant phenotype. Transcriptomics without the need for a reference genome is a powerful tool for comparative analyses given the availability of methods for data annotation and pattern identification. The strong resistance phenotype of MRI compared to SB22 likely led to the strength of the visible signal between the cultivars. NLRs that confer resistance to biotrophic pathogens have increased genetic diversity, and here we observe that, despite high genetic conservation between orthologs (Fig. 1), there are distinct genetic differences in the NLR repertoire (Van de Weyer *et al*., 2019).

Breeding cultivars with quantitative resistance has been shown to produce durable resistance, typically through the activity of multiple minor-effect genes (Brown, 2015; Niks *et al*., 2015). Other BDM resistant cultivars have been produced through interspecific hybridization of *O. basilicum* with *O. americanum* var. *pilosum*, leading to dominant resistance against two races of *P. belbahrii* (Ben-Naim & Weitman, 2021). We have observed that the downy mildew-resistant basil cultivars succumb to infection in production systems in different regions, consistent with the report of emerging pathogen races (Ben-Naim & Weitman, 2021). Therefore, understanding the physiological and molecular bases of host-pathogen interactions is critical to rapidly developing improved cultivars, monitoring strategies, and management practices. Identification of suitable molecular markers conferring multiple sources of resistance will improve breeding for basil downy mildew resistance and advance the understanding of molecular mechanisms of resistance and pathogenicity. These developments in approaches and knowledge can be broadly utilized by plant breeders and pathologists working with downy mildew pathogens on many different crops. After further validation, the genes predicted here can likely serve as molecular markers for the selection of downy mildew resistant sweet basils, and investigation and validation of physiological responses to infection may aid in developing more robust phenotyping assays. This comparative transcriptomics approach not only revealed candidate resistance genes, but also offered us new insights into differential infection responses in resistant and susceptible cultivars, which will open new avenues of investigation to further combat basil downy mildew.

This method of analysis is broadly applicable to two-organism biological systems where the identification of genes involved in any specific interaction is desired. Although genomes for holy basil (*O. tenuiflorum*) had been published prior to this study and sweet basil (*O. basilicum*) and *P. belbahrii* genomes have been published since, the methods described here worked exclusively from RNA sequencing data and did not require the whole genome sequence. Any system utilizing two organisms from different kingdoms could be examined using these methods as DNA sequences are differentiable down to reasonable taxonomic levels.

## Acknowledgements

We thank the MGHPCC for providing high-performance computing capacity for the RNAseq data analysis. This work was supported by the United States Department of Agriculture, Specialty Crop Research Initiative, National Institute of Food and Agriculture (USDA/SCRI/NIFA 2018-51181-28383) and USDA Hatch grant (MASR-2009-04374, MAS-2018-00496). L.-J.M. is also supported by an Investigator Award in Infectious Diseases and Pathogenesis by the Burroughs Wellcome Fund (BWF-1014893), the National Eye Institute of the National Institutes of Health (R01EY030150) and the National Science Foundation (IOS-1652641). We thank Chris Joyner and David O’Neil of the College of Natural Sciences Greenhouse at University of Massachusetts for continued support with plant propagation. We also thank Erin Patterson for assistance with phylogenetic analysis and Dr. Dilay Hazal Ayhan for help with allele expression analysis.

## Author Contribution

JS, L-JM, RP, and LG participated in the designing of the experiment. RP prepared all materials for the RNAseq, GAD performed all RNA-seq data analysis. KSA performed phylogenetic analysis and R-gene cloning, sequencing, and protein structural analyses. AG directed protein structural and sequence analysis. JM provided initial protein structural modeling analysis. KSA, GAD and L-JM wrote the manuscript. KSA and GAD prepared the figures, and all authors edited the paper.

## Disclaimer

This article was prepared while Anne Gershenson was employed at the University of Massachusetts Amherst. The opinions expressed in this article are the author’s own and do not reflect the view of the National Institutes of Health, the Department of Health and Human Services, or the United States government.

## Supporting Information

Additional supporting information may be found in the online version of this article.

**Fig. S1 Average Quality Scores.** Average quality scores plotted on the Y-axis by base number on the X-axis for all paired-end reads.

**Fig. S2 Pathogen Read Percentage.** Percentage of reads mapped to the *Peronospora belbahrii* sporangia data in MRI (black bars) and SB22 (grey bars) combined data at each timepoint plotted on the X-axis.

**Table S1** Primers Used In This Study

**Table S2** Coarse Clustering Results

**Table S3** Top ten structural homologs of MRI-R1 identified by DALI PDB Search

## Notes

### Competing Interest Statement

The authors have declared no competing interest.

### Summary of Updates

Supplemental files updated

