## Supplementary figures and images for "Identification of novel basil downy mildew resistance genes using *de novo* comparative transcriptomics"

### Supporting Information Figure S1

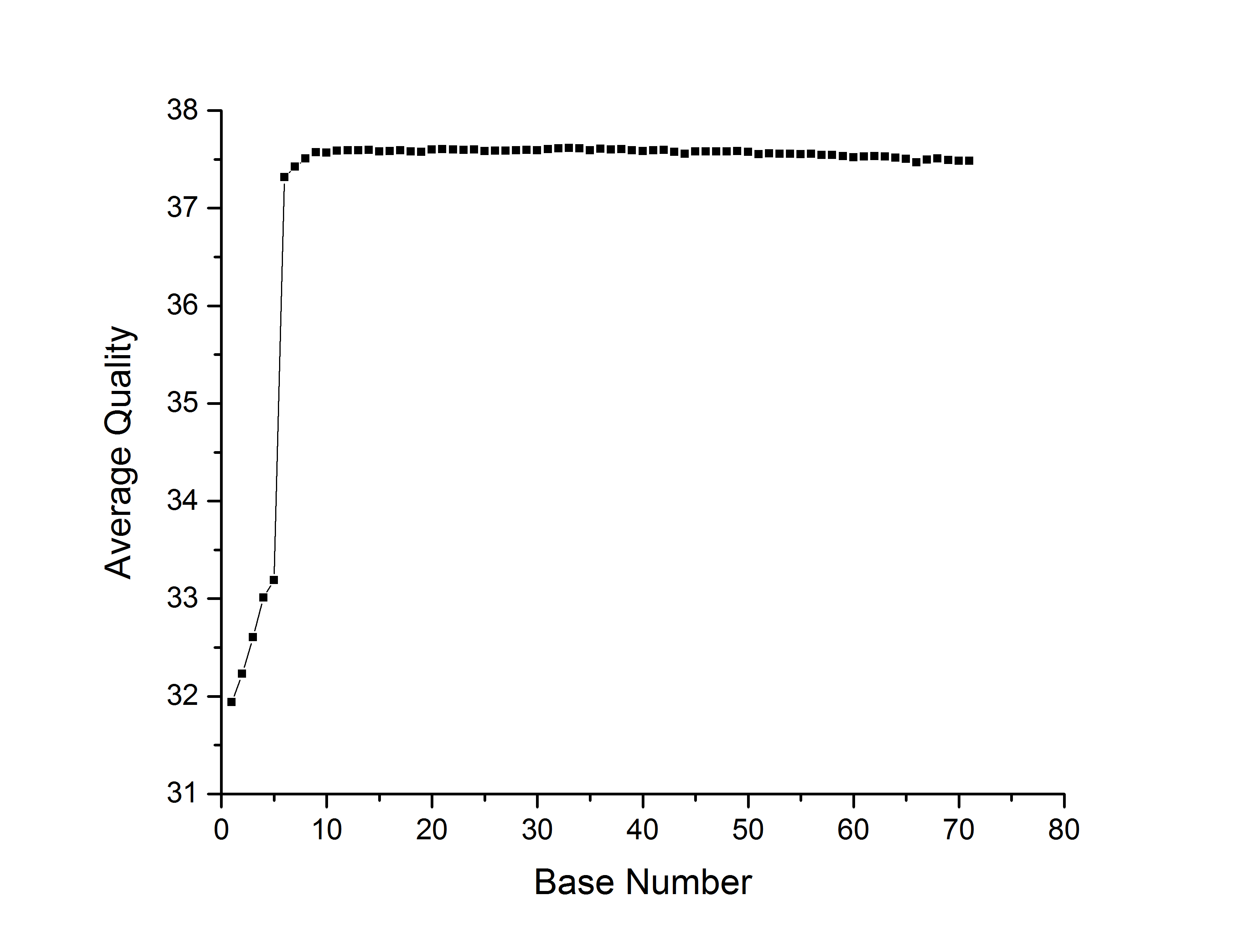

### Supporting Information Figure S2

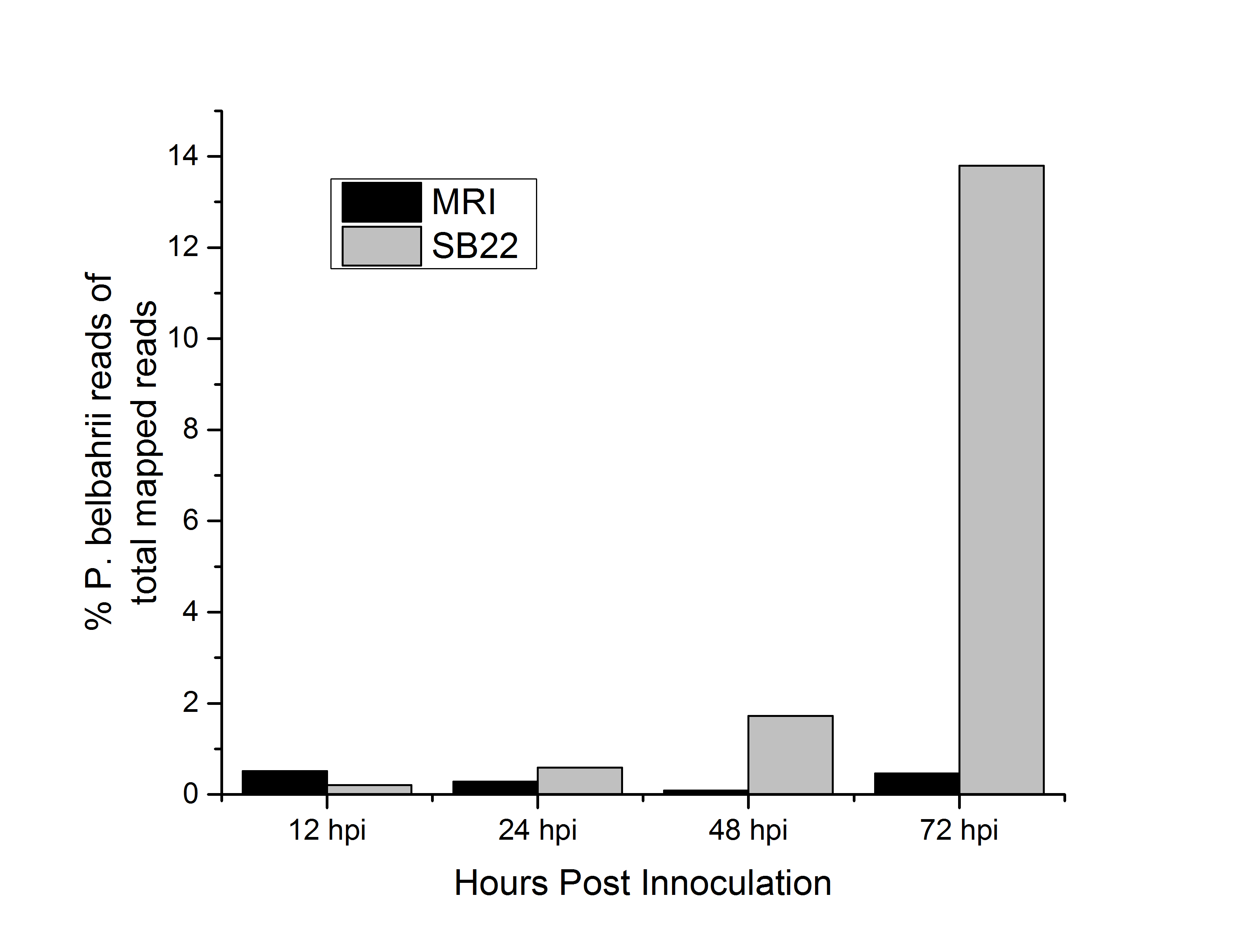
